# Correlative Fluorescence- and Electron Microscopy of Whole Breast Cancer Cells Reveals Different Distribution of ErbB2 Dependent on Underlying Actin

**DOI:** 10.1101/2020.01.14.906040

**Authors:** Indra Navina Dahmke, Patrick Trampert, Florian Weinberg, Zahra Mostajeran, Franziska Lautenschläger, Niels de Jonge

## Abstract

Epidermal growth factor receptor 2 (ErbB2) is found overexpressed in several cancers, such as gastric, and breast cancer, and is, therefore, an important therapeutic target. ErbB2 plays a central role in cancer cell invasiveness, and is associated with cytoskeletal reorganization. In order to study the spatial correlation of single ErbB2 proteins and actin filaments, we applied correlative fluorescence microscopy, and scanning transmission electron microscopy (STEM) to image specifically labeled SKBR3 breast cancer cells. The breast cancer cells were grown on microchips, transformed to express an actin-green fluorescent protein (GFP) fusion protein, and labeled with quantum dot (QD) nanoparticles attached to specific anti-ErbB2 Affibodies. Fluorescence microscopy was performed to identify cellular regions with spatially correlated actin and ErbB2 expression. For STEM of the intact plasma membrane of whole cells, the cells were fixed and covered with graphene. Spatial distribution patterns of ErbB2 in the actin rich ruffled membrane regions were examined, and compared to adjacent actin-low regions of the same cell, revealing an association of putative signaling active ErbB2 homodimers with actin-rich regions. ErbB2 homodimers were found absent from actin-low membrane regions, as well as after treatment of cells with Cytochalasin D, which breaks up larger actin filaments. In both latter data sets, a significant inter-label distance of 36 nm was identified, possibly indicating an indirect attachment to helical actin filaments via the formation of heterodimers of ErbB2 with epidermal growth factor receptor (EGFR). The possible attachment to actin filaments was further explored by identifying linear QD-chains in actin-rich regions, which also showed an inter-label distance of 36 nm.

## INTRODUCTION

The membrane protein ErbB2 also named HER2 is a member of the receptor tyrosine kinase family of epidermal growth factor receptors together with the epidermal growth factor receptor (EGFR, also named ErbB1), ErbB3, and ErbB4. ErbB2 plays a key role in cancer cell growth, and metastasis and, as such, resembles an important target for cancer treatment (Citri and Yarden, 2006; Henjes et al., 2012; Hynes and MacDonald, 2009). ErbB2 can homodimerize with another ErbB2 protein, or heterodimerize with other tyrosine kinases leading to the activation of the respective cell signaling pathways (Citri and Yarden, 2006). Overexpression of ErbB2 affects the actin cytoskeleton, and thus plays a role in the metastatic progression in cancer cells (Brix et al., 2014; Day et al., 2017), which resembles the major cause of mortality (Dillekas et al., 2019). The interaction of ErbB2 with actin was reported previously in several studies; for instance, activated ErbB2 was found to be spatially correlated with actin-rich membrane protrusions, like peripheral ruffles or filopodia-like protrusions, in ErbB2 overexpressing cells (Brandt et al., 1999; Chung et al., 2016; Jeong et al., 2016). HER2 homodimers were found to preferentially reside in membrane ruffles (Peckys et al., 2015).

In order to study the interplay of ErbB2 and actin filaments in membrane ruffles, we applied correlative fluorescence microscopy and liquid phase scanning transmission electron microscopy (STEM) of whole cancer cells. SKBR3 breast cancer cells were grown on microchips, transformed to express an actin-green fluorescent protein (GFP) fusion protein, and ErbB2 was labeled via an Affibody in a two-step procedure with a quantum dot (QD) nanoparticle. The cells were covered with a sheet of graphene to prevent drying out in the vacuum of the electron microscope (Dahmke et al., 2017). By analysis of ErbB2 receptors at the single-molecule level using STEM in cell membrane regions of varying actin content, we observed different distribution patterns of membranous ErbB2 in dependency of the underlying actin filaments.

## MATERIALS AND METHODS

### Cell culturing and labeling of Actin in SKBR3 breast cancer cells

Human breast cancer SKBR3 cells (ATCC^®^ LGC Standards GmbH, Germany, HTB-30^™^) were cultured in 25 cm^2^ cell culture flasks (Greiner Bio-One GmbH, Germany, Cellstar^®^, TC^™^) with Dulbecco’s Modified Eagle’s Medium GlutaMAX^™^ (high glucose and pyruvate, DMEM, Gibco, Thermo Fisher Scientific, USA) containing 1% non-essential amino acids and 10% heat inactivated Fetal Bovine Serum (FBS, Gibco, Thermo Fisher Scientific) under standard cell culturing conditions (at 37 °C and 5% CO_2_). SKBR3 cells serve as a common model system for ErbB2-overexpressing breast cancer (Henjes et al., 2012).

For correlative fluorescence and electron microscopy, cells were seeded two days prior to imaging on custom-made SiN-microchips (dimensions: 2.0 × 2.6 × 0.3 mm^3^, SiN-window: 0.50 × 0.15 mm^2^ of 50 nm thickness, DensSolutions, The Netherlands) (Ring et al., 2011). In preparation, the microchips were cleaned with ArO_2_-plasma for 5 min, then coated with poly-L-lysine (0.01%, Sigma-Aldrich, Germany) for 5 min at room temperature and washed twice with phosphate buffered solution (1×, pH 7.4, Gibco, Thermo Fisher Scientific). The microchips were then coated with fibronectin (15 μg/mL, Sigma-Aldrich) under the same conditions. Immediately after, the microchips were transferred to a 96-well-plate (Greiner Bio-One GmbH, Cellstar^®^) and covered with 100 μL of FBS-free DMEM each. Next, SKBR3 cells were harvested with CellStripper (Corning, USA) and diluted to 200,000 cells per mL with 100 μL of the prepared cell-suspension added to each well. After 2-3 h of incubation under standard culture conditions, the microchips were transferred to new wells pre-filled with 200 μL of DMEM with CellLight^™^ Actin-GFP, BacMam 2.0 (30 particles per cell, Invitrogen, Thermo Fisher Scientific) to transduce the cells with actin-GFP. The samples were cultivated for 40 h and either treated with an actin-inhibitor (see below) for 1h or immediately labeled with anti-ErbB2-Affibodies.

### Labeling of ErbB2

For the labeling of ErbB2 molecules with quantum dots (QDs), microchips were rinsed once in GS-BSA-PBS (1% goat-serum (GS), Gibco; 1% BSA (molecular biology-grade albumin fraction V, Carl Roth GmbH-Co. KG, Germany); in PBS) and incubated in the same solution for 5 min at 37 °C to block unspecific binding sites before incubation with the biotin-conjugated anti-ErbB2-Affibodies^®^ ((ZERBB2: 342)2, ErbB2-AFF-B, Affibody, Sweden). Next, microchips were incubated for 10 min at 37 °C with 200 nM ErbB2-AFF-B in GS-BSA-PBS. After consecutive washing with 1% BSA-PBS (2×), PBS(1×) and with cacodylate-buffer (CB, 0.1 M sodium cacodylate trihydrate, Carl Roth GmbH, Germany, and 0.1 M saccharose, pH 7.4), the cells were fixed for 15 min at room temperature with 4% formaldehyde (Electron Microscopy Sciences, USA) in CB to prevent QD-induced clustering of ErbB2-molecules. Next, cells were rinsed once with CB and three times with PBS-BSA, then incubated in 0.1 M glycine in PBS for 2 min. After two additional washes with PBS, cells were incubated with streptavidin-conjugated Qdot 655 (5 nM, Life Technologies, USA) in borate buffer (40 mM, sodium tetraborate boric acid, Sigma-Aldrich, pH 8.3) at room temperature for 12 min. Finally, cells were washed three times in PBS-BSA (1%) and subjected to fluorescence microscopy. Further details of the protocol, including control experiments to test for unspecific binding of QDs, are reported elsewhere (Peckys and de Jonge, 2015; Peckys et al., 2015).

After fluorescence microscopy of the samples, an additional fixation step was added to enhance the stability of the cellular structures for imaging by means of electron microscopy (EM). The cells were fixed with 2% glutaraldehyde in CB buffer for 12 min, and rinsed once with CB and three times with PBS-BSA. Samples were stored in PBS-BSA at 4 °C until later coverage with graphene.

### Disruption of actin containing ruffles

Cytochalasin D (Sigma-Aldrich, Germany) was applied to SKBR3 cells to disrupt the formation of peripheral ruffles by disturbance of the actin network. For this purpose, the cells were transduced with actin-GFP and seeded on Si microchips as described above. After 40 h of cultivation, cells were examined by light microscopy prior to incubation with either DMSO (negative control, Carl Roth GmbH-Co. KG, Germany) or 2 μM of Cytochalasin D in GS-BSA-PBS for 60 min at 37 °C and 5% CO_2_. This concentration has been applied to SKBR3 cells (Stofega et al., 2004) before and shown inhibition of membrane ruffling after 10 min but no ultrastructural alterations in the microfilament bundles (Yahara et al., 1982). Incubation with the anti-ErbB2-Affibody for 10 min at 37 °C and 5% CO_2_ followed, and a staining procedure was performed as described above.

### Light microscopy

After fixation with formaldehyde, the samples were imaged with an inverted light microscope (DMI 6000B, Leica, Germany). The microchips were placed upside-down in a plasma-cleaned, glass bottom cell culture dish (35 mm in diameter, MatTek Coperation, USA) filled with PBS-BSA (1%) and imaged with 20× (HCX PL Fluo TAR L, 0.40, dry, Leica), 40× (HCX PL Fluo TAR L, 0.60, dry, Leica), or 63x oil (HC PL APO 63x/1.40 oil) objectives in direct interference contrast (DIC) and fluorescence mode (filter cube L5: excitation: 460-500 nm, DIM: 505 nm, emission: 512-542 nm and TX2: excitation: 540-580 nm, DIM: 595 nm, emission: 607-683). Cellular regions expressing high levels of ErbB2 emitted a red fluorescence at λ=655 nm, whereas actin fibers emitted green fluorescence at λ=520 nm. The images were acquired by employing the LAS FX operational software version 3.4.1.17822 (Leica) and manually stitched together to create overview images.

### Graphene deposition

Polymethyl methacrylate (PMMA) covered graphene (“trivial transfer graphene”, ACS Materials, USA) was removed from its polymer support according to instructions of the manufacturer and left overnight in a 10% sodium persulfate solution. This procedure removed potential copper contaminants from the production process, which interfere with protein labeling for STEM. Afterwards, the graphene film was washed in deionized H2O three times, transferred to a salt crystal (International crystal laboratory, USA), and baked for 20 min at 100°C. Finally, the PMMA was removed from the graphene film by incubating the graphene-salt in acetone (100%, high pressure liquid chromatography grade, Carl Roth GmbH-Co. KG) for 30 min at 30 °C. This step was repeated two times. To transfer the graphene film onto cancer cells grown on microchips, the graphene-NaCl crystals were placed on the surface of pure H2O (Carl Roth GmbH-Co. KG), as described elsewhere (Dahmke et al., 2017; Textor and de Jonge, 2018). Once the underlying NaCl was completely dissolved, the free-floating graphene film was positioned over the microchip and lifted up. The position of the graphene film over the SiN-window was then verified with a binocular. The microchip was left to air dry at room temperature for about 5 min, then stored in a humid chamber at 4 °C until imaging.

### STEM

STEM images were acquired with a transmission electron microscope (TEM) (JEM-ARM 200F, JEOL, Japan) equipped with a cold field emission gun, and a STEM probe corrector (CEOS GmbH, Germany). The microchip was mounted on a standard TEM sample holder (JEOL). The electron beam energy was 200 keV. In STEM mode, images were recorded with a pixel dwell time of *τ* = 14 μs. The electron probe setting was 4c, combined with an aperture CL2-3 with a 20 μm diameter, resulting in a probe current of *I_p_* = 63 pA, and an electron dose of 67 e^−^/Å^2^ for a magnification of 100,000× (the pixel size was 1 nm with 2048 x 2048 image resolution). The camera length was set to 8 cm, leading to a detector opening semi-angle of 43 mrad for the angular dark field detector. This relates to the opening through which electrons pass the detector, although, the actual collection angle is larger.

### QD-label analysis

For the detection of QD-labeled ErbB2 membrane proteins in the STEM micrographs, an automated analysis programmed in ImageJ (National Institute of Health, USA) was applied (Peckys et al., 2015). To avoid false counting of label positions, STEM images were preprocessed by checking for contaminating, electron-dense particles different from QDs that are of a large size. In case contaminating particles were present, STEM micrographs were altered by manually greying out the latter contaminations. The images were further processed automatically. A Gaussian filter was first applied with a radius of 1.5 pixel for noise-filtering. Variations in the image background were then filtered using a Fourier filter. The image was subsequently binarized by applying an automated threshold with maximal entropy settings. The program accounted for a bin width set to 2 nm and particles with a radius > 10 nm. For the subsequent statistical analysis using a pair correlation function g(r), a locally designed software tool in C++ was used (Peckys et al., 2015). The pair correlation function is defined as (Stoyan and Stoyan, 1996):

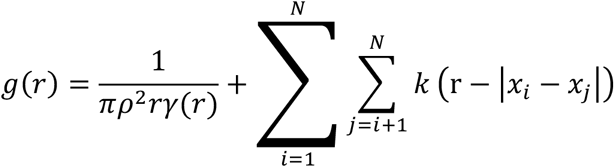

with *r*, the radial distance, and *ρ*, the labeling density in the image. The covariance function, *γ*(*r*), and the kernel, *k*, are defined elsewhere (Stoyan et al., 1993). The modulus, |*x_i_ − x_j_*|, characterizes the distance between two reference points, *i* and *j*, with *x* indicating the twodimensional coordinates, (*x, y*), of particles in the analyzed micrograph. This calculation of *g*(*r*) assumes a planar location of the QD-labeled membrane proteins. The QD-label distribution was analyzed for different experimental groups by adding all label positions acquired in all images of that group.

In order to test the significance of the results of the statistical analysis by the pair correlation function, software of local design in c++ was applied to generate data sets with a random particle distribution, but a definable particle density. The diameter of the particles was 12 nm and overlap of particles was not allowed. For comparison of the experimental data derived from the STEM images, the same number of random images as in the respective experimental data set was generated to contain the average QD-particle density of this set. The generated random data was then subjected to pair correlation analysis in the same way as the experimental data sets.

### QD-particle chain analysis

Linear QD-particle chains were automatically searched in the STEM images (=experimental datasets). As shown before in experimentation (Parker et al., 2018), six particles in order suffice for building such a particle chain. This was experimentally validated with the help of a statistical analysis. The same parameters as described before were selected meaning (i) the distance between consecutive particles on a line was allowed to be 36 nm ± 15 nm, and (ii) the angle of connecting lines between three particles in a row was allowed to be 180° ± 30°. The experiments were conducted in MatLab (MATLAB and Statistics Toolbox Release 2018a. The MathWorks, Inc., Natick, Massachusetts, USA), using an in-house developed program (Parker et al., 2018). The corresponding pseudo-code is illustrated in **Scheme 1**.

Additionally, random particle data was generated and evaluated. Each investigated image was divided into 8 × 8 segments, and then particle positions in each segment were randomized. This procedure ensures that the large-scale structures in the image, for example localized protein densities, are preserved, but exact positioning is discarded. The algorithm was applied to the randomized data and the results were compared to the linear structures detected in the experimental data, in order to show that our observations were not based on random effects.

## RESULTS

### Method for single-molecule microscopy of ErbB2 correlated to presence of actin

A correlative microscopy method was established to analyze the spatial correlation of the functional state of ErbB2 with the abundance of actin in whole cancer cells (**Figure 1**). SKBR3 breast cancer cells grown on silicon microchips were first transduced with an actin-GFP baculovirus construct, and cultivated for 40 h. Optionally, breast cancer cells were incubated with Cytochalasin D for 1 h to induce de-polymerization of the actin cytoskeleton and disruption of membrane ruffles (Schliwa, 1982). In the following steps, the breast cancer cells were incubated with a specific biotin-conjugated anti-ErbB2-Affibody, fixated with paraformaldehyde to prevent artificial clustering of labels, and marked with fluorescent, streptavidin-conjugated QDs. Cell regions of interest containing higher amounts of both actin and ErbB2 were then identified using Fluorescence microscopy (**Figure 2**).

**Figure 1.**
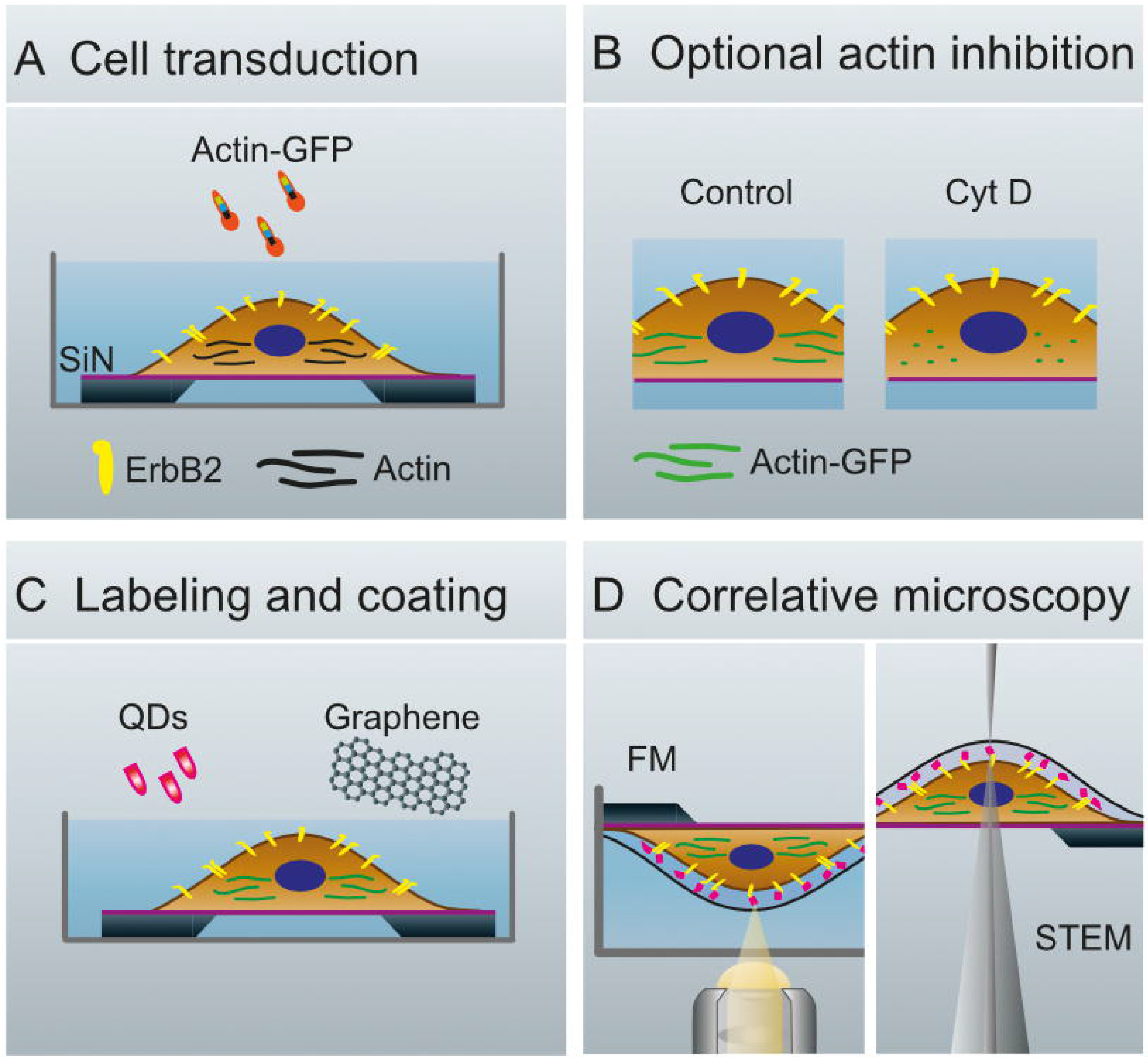
Schematics of whole cell preparation for the analysis of membrane proteins. **(A)** Transduction of SKBR3 cells, seeded on silicon microchips with a silicon nitride membrane, with actin-green fluorescent protein (GFP)-constructs and cultivation of cells for 40 h. (**B**) Cells remain untreated (Control, left) or are treated with Cytochalasin (Cyt) D (right). **(C)** Membranous ErbB2 is labeled with streptavidin coated quantum dots (QDs) via Affibody-biotin conjugates. The cells are covered with graphene to protect against evaporation of the liquid in the vacuum of the electron microscope. (**D**) Configuration for fluorescence microscopy (FM, left), and for scanning transmission electron microscopy (STEM, right).

To protect the hydrated samples from drying when placed in vacuum during EM, each microchip was covered with a multi-layered graphene film, using a method of transferring graphene floating on a water surface onto the sample (Figure 1) (Dahmke et al., 2017). The samples were then imaged with STEM, first at low magnification (1000-2000 ×) in order to correlate the cell locations in the images obtained by Fluorescence microscopy with the ones obtained by means of EM (**Figure 2A-D**). Areas of interest as determined by Fluorescence microscopy were identified in the EM images, and examined using STEM at higher magnification (100,000-120,000 ×) so that single labels were visible (**Figure 2E, F**). From this, the functional state of ErbB2 was determined as described in the following section.

**Figure 2.**
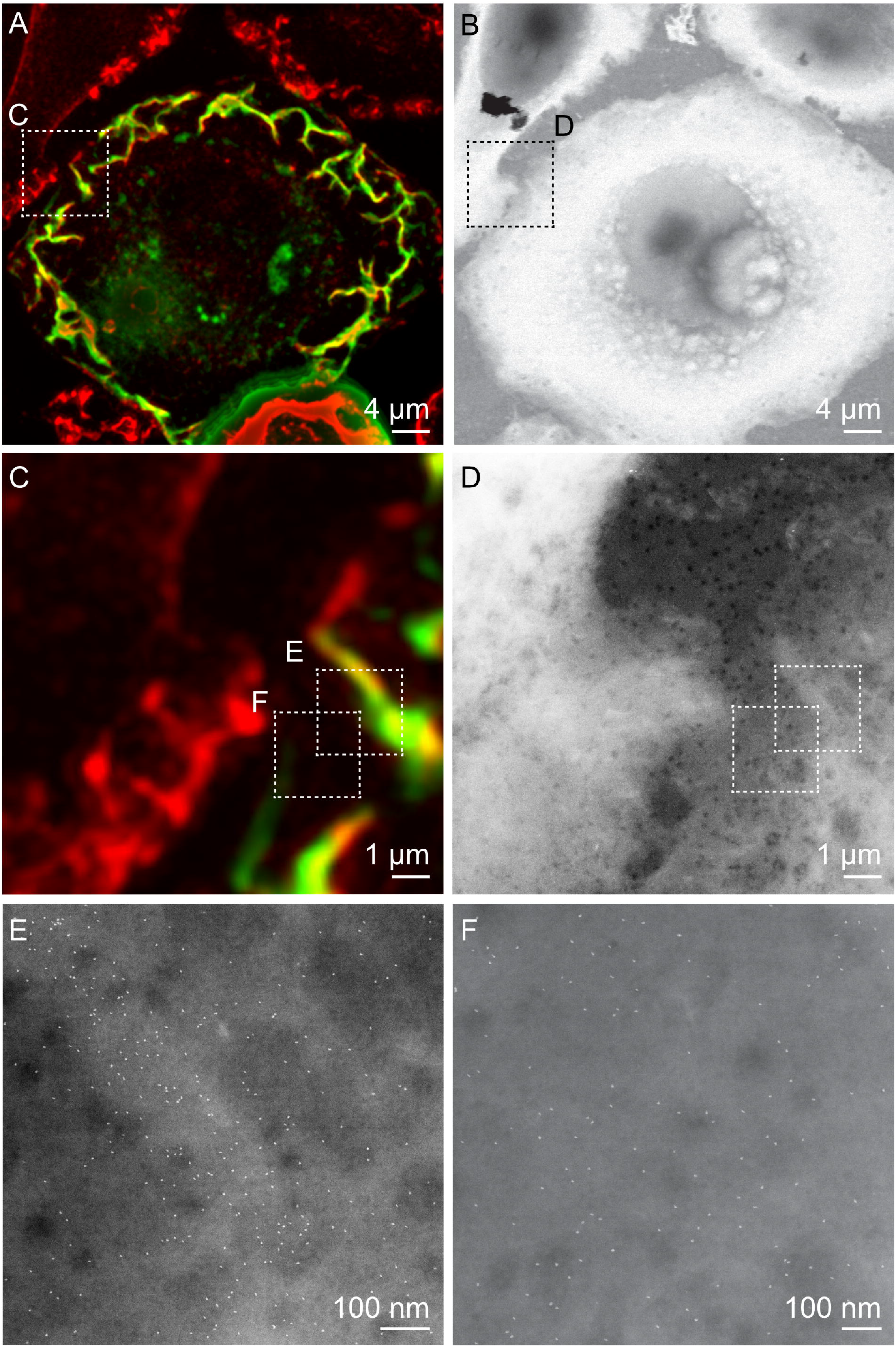
Correlative fluorescence microscopy and STEM of whole breast cancer cells. **(A, C)** Cropped fluorescence micrographs of SKBR3 breast cancer cells showing cellular actin-GFP in green and QD-labeled membrane ErbB2 in red. Areas where both signals overlap appear yellow. Actin-containing ruffles can be identified as bright green and yellowish lines at the cell edges. **(B, D)** Corresponding STEM micrographs of graphene-covered breast cancer cells acquired at the same spots. B: Magnification *M* = 1,000×, D: *M* = 4,000×. **(E, F)** STEM micrographs of regions marked in C, and D acquired at higher magnification than in D. QD-labels appear as white dots. Membrane ruffle in E appears in a light grey shade compared to flat cell regions in F, appearing in a darker grey, E: *M* = 100,000×, F: *M* = 120,000×.

### Statistical method to determine functional state of ErbB2

In order to distinguish ErbB2 proteins forming homodimers from ones that were randomly grouped at closer vicinities, the label position data was statistically analyzed using the pair correlation function, *g*(*r*) (Peckys et al., 2015; Stoyan et al., 1993). This function, gives the probability of finding two labels at a certain radial distance *r* with *g*(*r*) = 1, indicating random positioning. The expected average center-to-center distance of 20 nm between two QDs accounts for each attaching to an ErbB2 protein in a homodimer (Peckys et al., 2015); however, this value has some variation due to flexibility of the linkers. The functional state of ErbB2 is thus detected via labeling of single, endogenous ErbB2 proteins with a high abundance in the cell membrane (Yang et al., 2007). Such data is not obtained with other techniques, for example, using whole antibody complexes or artificial fusion proteins as labels (Liu et al., 2007; Needham et al., 2013). The subsequent statistical analysis corrects for false-positive readings by excluding randomly paired labels; this may be the case with proximity techniques, such as proximity ligation or fluorescence resonance energy transfer assays (Greenwood et al., 2015; Needham et al., 2013).

### Distribution of membranous ErbBs in relation to actin-GFP

In order to correlate the distribution of QD-labels with the occurrence of actin-GFP, STEM images were analyzed as collected at two different membrane regions. For the first group, STEM images were collected in cell areas identified by the presence of the fluorescence signal for actin-GFP and peripheral membrane ruffles identified in DIC mode (**Supplementary Figure S1**), as described elsewhere (Peckys et al., 2015). Here, we found a higher density of QD-labels compared to cell areas with a low content of actin (**Table 1**). The resulting graph of *g*(*r*), collected at cell actin rich cell regions, displayed a peak at 20 nm, which exceeds random levels (**Figure 3A**). This peak is significantly above the statistical fluctuations, since *g*(*r*)=1.7 - 1 > 3\**SD*, with the standard deviation, *SD* being 0.02 of the *g*(*r*) for 50 nm < *r* < 200 nm in this case, and the factor of 3 originating from the Rose criterion (Rose, 1973). This suggests the plasma membrane area contained signaling-active homodimers (Citri and Yarden, 2006; Peckys et al., 2015). For comparison, a simulation of a random distribution of QD-labels of this size was included in **Figure 3A** as well, showing a similar level of statistical fluctuations as the experimental data but the absence of a peak at 20 nm

**Figure 3.**
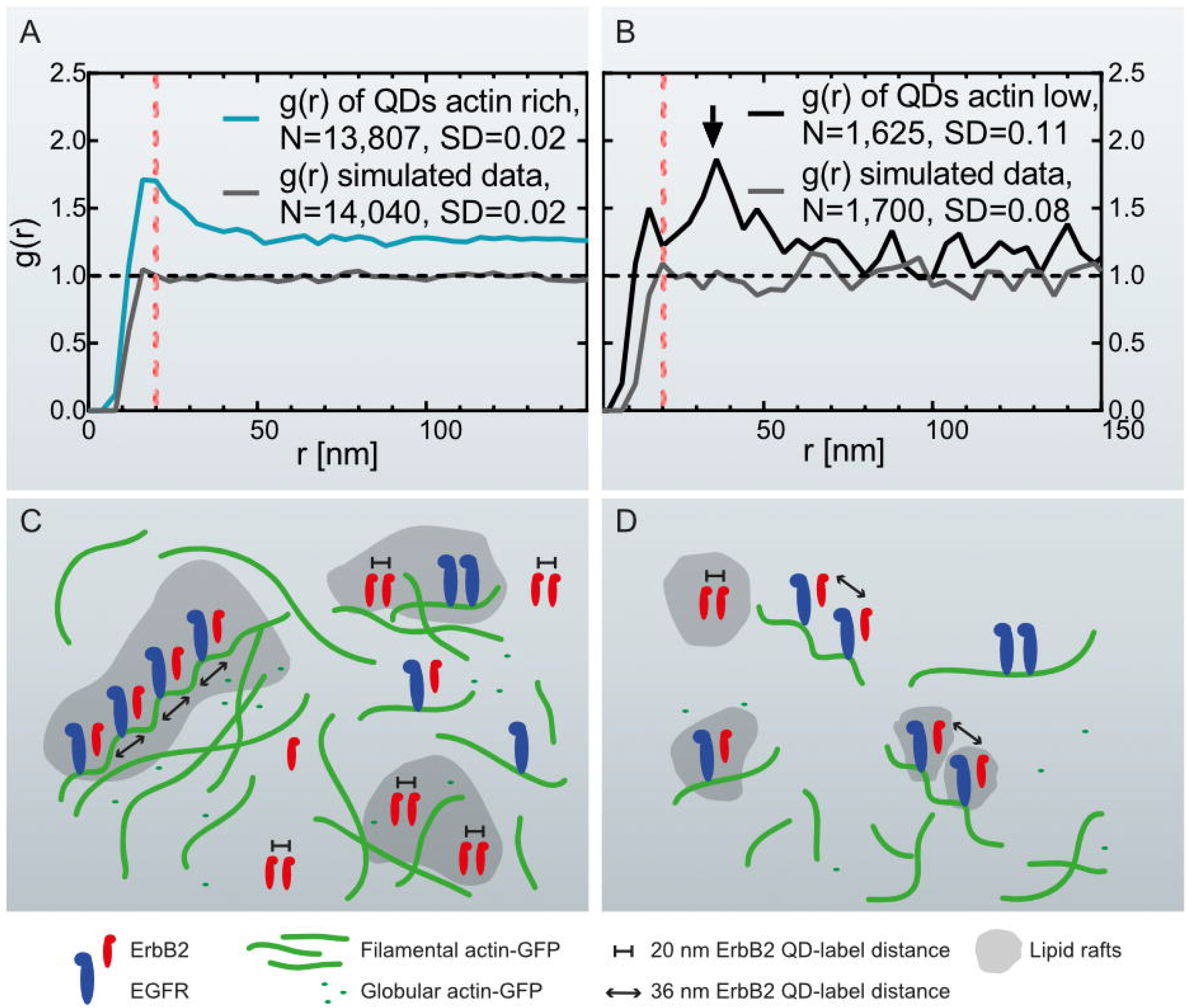
Graphs of the pair correlation function *g*(*r*) versus the radial distance *r* between QD-labels, collected at actin-rich- and actin-low cell regions with correlating schemes. **(A)** *g*(*r*) of QD-labels detected in actin-rich cellular regions collected from 27 images of 11 cells. The red dotted line marks *g*(*r*) = 20 nm. As comparison, *g*(*r*) of simulated data is included of randomly positioned labels with the same particle density (see **Table 1**). **(B)** *g*(*r*) of QD-labels in actin-low regions collected in 17 images of 8 cells. The arrow marks a peak at *r* = 36 nm. The red dotted line is at *r* = 20 nm. *g*(*r*) of simulated data of randomly positioned labels is also included. **(C)** Schematic representation of the assumed distribution of ErbB2 and EGFR homo- and heterodimers in relation to cortical actin filaments and lipid rafts in actin-rich, ruffled cell regions. EGFR is bound to actin filaments while ErbB2 can move freely in the membrane and lipid rafts. Those ErbB2 molecules assembled in ErbB2:EGFR heterodimers are bound to helical actin filaments with a pitch of 36 nm. The number of ErbB2 homodimers in ruffled regions outweighs the number of postulated heterodimers. (**D**) Actin-low, flat cell regions with fewer ErbB2 homodimers being present than in ruffled regions, and postulated heterodimers dominating the analysis. The ruffled regions contain a higher number of growth factor receptors than the flat regions.

**Table 1.** Summary of scanning transmission electron microscopy (STEM) data used for label distribution analyses. STEM images were acquired in three different experiments to compare the distribution of quantum dot (QD)-labels by pair correlation analysis. Data sets comprised of STEM images acquired at actin-rich, actin-low regions, and Cytochalasin D (CytoD) treated SKBR3 breast cancer cells that were transduced with actin-green fluorescent protein (GFP).

| Experimental group | No. experiments | No. of cells | No. of images | No. of all particles | Total area ($\mu\text{m}^2$) | Average particle density per image (particles/$\mu\text{m}^2$) |
| --- | --- | --- | --- | --- | --- | --- |
| Actin low | 3 | 8 | 17 | 1,640 | 70 | 24 |
| Actin rich | 3 | 11 | 27 | 13,807 | 110 | 124 |
| Cyto D | 2 | 5 | 7 | 400 | 25 | 16 |

For the second group, STEM micrographs were collected in adjacent membrane regions with lower signal for actin-GFP as compared to the first group. In contrast to the *g*(*r*) of actin-rich regions, the *g*(*r*) of actin-low membrane areas showed a peak at 36 nm, rather than at 20 nm (**Figure 3B**). This graph also displays larger fluctuations due to the lower number of collected QD positions compared to the actin-rich regions (**Table 1**) as a result of a smaller number of detected labels. A similar level of fluctuations is visible in the corresponding random simulation of QD positions of the same density and amount (**Figure 3B**). The peak at 36 nm is nevertheless significant, since the *SD* of the *g*(*r*) is 0.11 (for 50 nm < *r* < 200 nm) and the peak at since *g*(*r*)=1.8 - 1 > 3\**SD*.

The presence of a 36 nm peak was possibly caused by a substantial fraction of ErbB2 interacting with overexpressed actin in the analyzed cells, since it was previously hypothesized that ErbB2 would bind in a chain-like orientation to actin (Parker et al., 2018) at intervals of ~36 nm (**Figure 3C, D**) (Needham et al., 2014). To test if the peak was caused by a high expression level of actin, a pair correlation analysis was performed of QD-labels in flat membrane regions of cells that were negative for actin-GFP in the fluorescence images. In accordance with former findings, no peak at 36 nm was detected (**Supplementary Figure S2**). A further observation for the largest range of *r* in all experiments is that *g*(*r*) > 1. This observation can be either interpreted as clustering of groups of labels or as the rising curvature of the plasma membrane of the cancer cells resulting in *g*(*r*) > 1 for the inter-label distances (Peckys et al., 2019).

### Cytochalasin D induces changes on membranous ErbB2 distribution

Next, the formation of ErbB2 homodimers was tested at peripheral membrane ruffles to test dependence on the underlying actin network. SKBR3 cells were treated with 2 μM Cytochalasin D for 1 h to disturb membrane ruffling. Cytochalasin D is a fungal metabolite that binds to the barbed end of actin filaments and inhibits the association and dissociation of actin monomers (Cooper, 1987). This treatment causes a disruption of the actin network, and induces the formation of actin foci, cell rounding, and inhibition of peripheral ruffle formation (Schliwa, 1982; Yahara et al., 1982). **Supplementary Figure S3** shows that Cytochalasin D induces loss of membrane ruffling, and cell rounding in SKBR3 cells. Furthermore, Cytochalasin D incubation induced the formation of distinct fingerlike protrusions, known as dendrites (Ting-Beall et al., 1995), which can be observed in the STEM image of Cytochalasin D treated cells (**Figure 4A, B**). QD-labels in these affected membrane regions gave a surface density lower than that measured in cell regions with high and low actin contents in untreated cells (**Table 1**). Subsequent statistical analysis of the QD-labeled ErbB2 proteins showed an ErbB2-pattern similar to that of flat areas in untreated cells where the peak at 20 nm was lost, but a prominent peak at 32 - 36 nm was registered (**Figure 4C**). The resulting peak at *g*(*r*)=3.9 is significant, since the *SD* of the *g*(*r*) is 0.42 (for 50 nm < *r* < 200 nm) and the criterion *g*(*r*)= - 1 > 3\**SD* applies. These results confirm that the formation of ErbB2 homodimers is coupled to an intact actin network in the membrane of SKBR3 cells.

**Figure 4.**
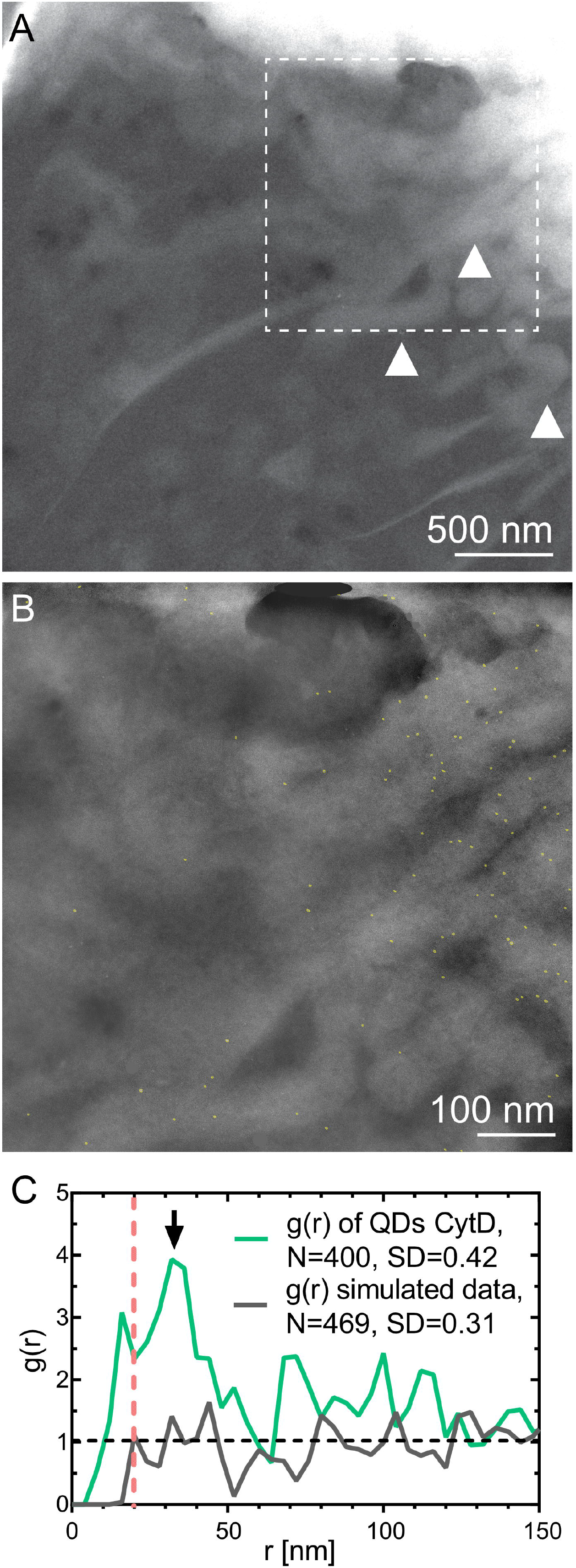
STEM of whole breast cancer cells treated with Cytochalasin D. **(A)** STEM of graphene-covered SKBR3 breast cancer cells (*M* = 30,000×). Cytochalasin D induced-fingerlike protrusions are marked exemplarily with white arrowheads. **(B)** STEM image of region enclosed with dashed square in A (*M* =100,000×). Automatically detected QD-labels are marked in yellow to enhance visibility. **(C)** *g*(*r*) of QD-labels collected in 7 images of cell edges of Cytochalasin D treated cells. The arrow marks a peak at *r* = 36 nm, and the dotted line indicates *r* = 20 nm. *g*(*r*) of simulated data of randomly positioned labels.

### Detection of linear QD-chains in actin rich membrane regions

After the detection of the inter-label distance of 36 nm above random in certain experimental data sets, a search for linearly arranged QDs was conducted in the STEM images of all data sets, namely, actin-GFP-rich, actin-GFP-low, and Cytochalasin D treated cells (Parker et al., 2018). A linear particle chain was defined to consist of six consecutive QDs with an interlabel distance of 36 nm ± 15 nm, and an angle of three consecutive QDs in a row of 180° ± 30°. Linear chains complying with these parameters were detected in three STEM images acquired in actin-GFP-rich regions, of which one example is given in Figure 5, but not in the other datasets (**Table 2**). To test if the identified lines were caused by false-positive pattern-detection, random particle data was generated for the according particle densities. For each image, 100 randomized data sets were generated, and the probability of finding at least the number of particle-chains that were detected in the according experimental data set (= image) was calculated. The calculated *p*-values were below 0.04 for all three data sets, indicating that the detected QD-particle chains did not form randomly. Thus, it was deduced that a fraction of ErbB2 interacted with filaments of the actin cortex in actin-GFP rich region.

**Table 2.**
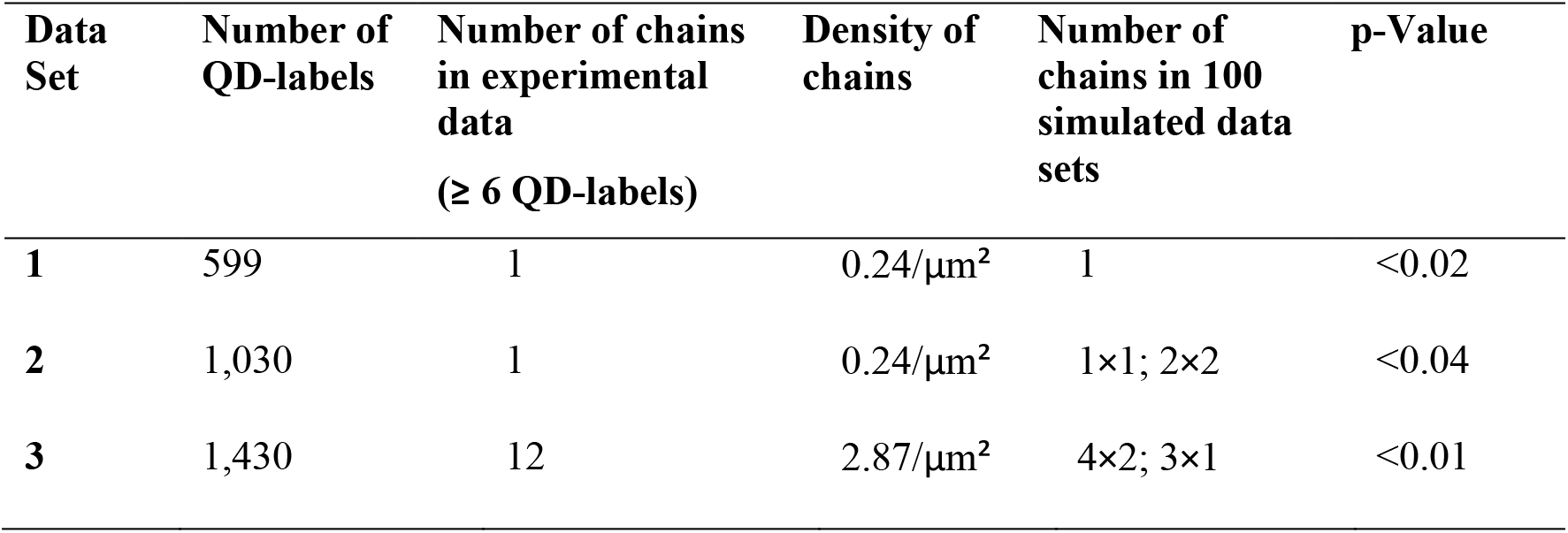
Comparison of experimental and simulated random data of linear QD-chains on cancer cells. Linear chains of QD-labels of ≥ 6 QD-labels with an inter-label distance of 36 nm ± 15 nm and angle of 180° ± 30° were detected in a number of experimental STEM images. For each experimental data set (one image), 100 randomly simulated data sets were generated with the same label density as found in the experimental data. The number of chains in the simulated data reflects the number of randomly simulated data sets in which one or two chains per data set were found. The probability of finding at least the same number of chains detected in the experimental data in the simulated data is listed in the last column. A p-valued < 0.05 was considered significant.

| Data Set | Number of QD-labels | Number of chains in experimental data ($\geq 6$ QD-labels) | Density of chains | Number of chains in 100 simulated data sets | p-Value |
| --- | --- | --- | --- | --- | --- |
| 1 | 599 | 1 | 0.24/$\mu\text{m}^2$ | 1 | <0.02 |
| 2 | 1,030 | 1 | 0.24/$\mu\text{m}^2$ | 1×1; 2×2 | <0.04 |
| 3 | 1,430 | 12 | 2.87/$\mu\text{m}^2$ | 4×2; 3×1 | <0.01 |

**Figure 5.**
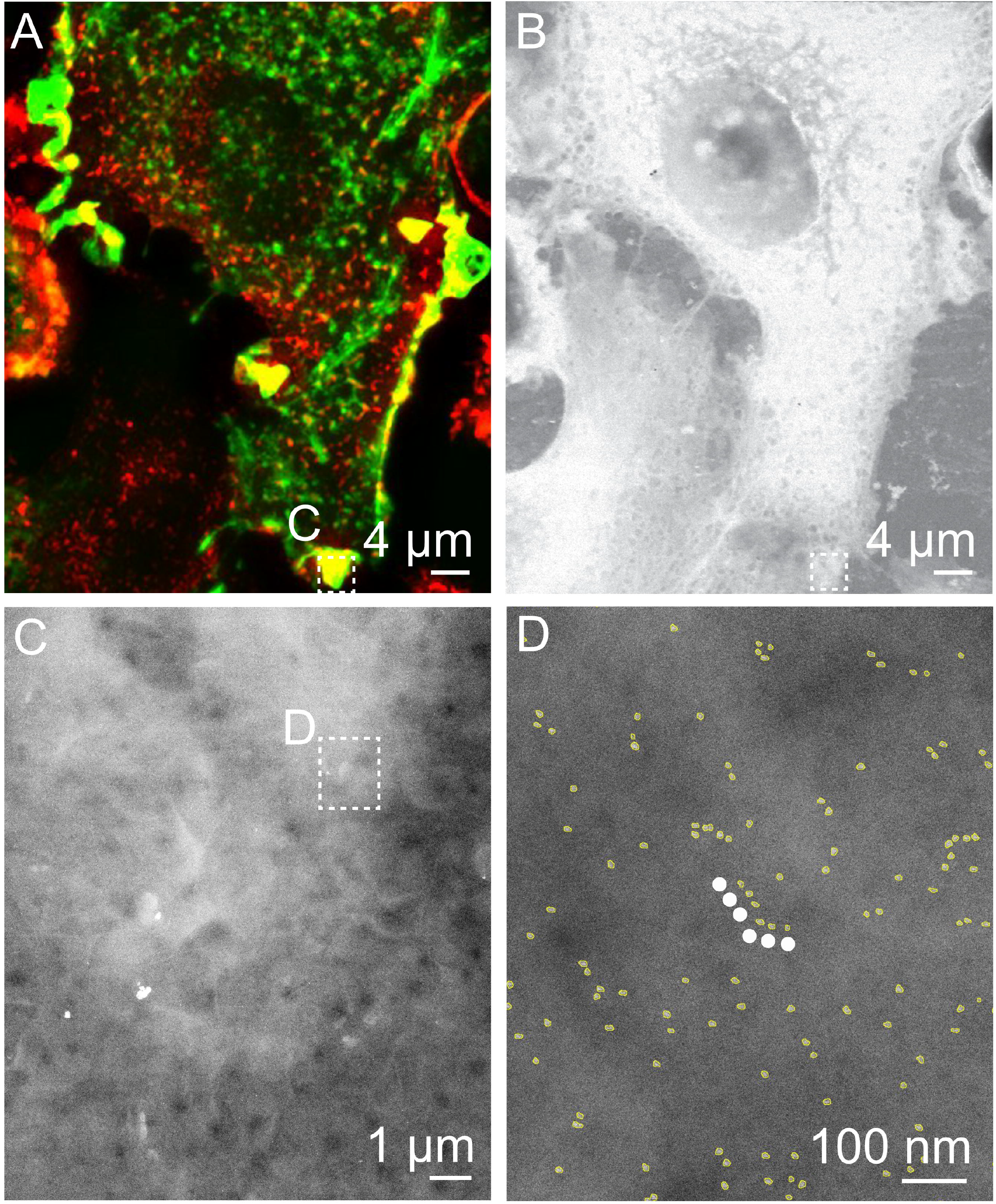
Correlative fluorescence microscopy and STEM of QD-labeled ErbB2 on whole breast cancer cells revealing linear QD-chains in actin rich region. **(A)** Cropped fluorescence micrograph of a SKBR3 breast cancer cell showing cellular actin-GFP in green and QD-labeled membrane ErbB2 in red. Areas where both signals overlap appear yellow. **(B)** Corresponding STEM micrograph acquired at the same spot (*M* = 1,000×). **(C)** STEM micrograph of region marked in A. B at *M* =30,000×. **(D)** Region selected from C imaged at *M* =100,000 ×. Automatically detected QD-labels are marked in yellow to enhance the visibility. The white dotted line marks a linear QD-chain consisting of six QD-labels with an inter-label distance of 36 nm ± 15 nm and angle of 180° ± 30°.

## DISCUSSION

In this study, we applied correlative Fluorescence microscopy and STEM of graphene-covered whole cancer cells to investigate distribution patterns of membrane ErbB2 in spatial relation to actin. Previously, ErbB2 homodimers were observed residing preferentially in ruffled cellular regions (Peckys et al., 2015). This prompted the question of whether ErbB2 proteins were coupled to actin as a mechanism explaining its preferential location in these membrane areas. Our new data now indicates that a spatial linkage between actin and ErbB2 exists. Our microscopy approach facilitated the detection and quantification of endogenous ErbB2 membrane proteins with a high abundance at the single molecule level. This allowed for the assessment of the functional state of ErbB2 proteins, as well as for the spatial correlation with underlying structures. Correlative light- and electron microscopy of whole cells in hydrated state (Peckys et al., 2015; Sato et al., 2017) is advantageous for studying protein function compared to conventional light microscopy, flow cytometry or gel electrophoresis, which provide lower spatial resolution or lack the possibility of spatial analysis altogether (Peckys et al., 2015). A subsequent statistical analysis of our data was performed to verify the presence of ErbB2 homodimers, and excluding randomly positioned protein pairs from the analysis, which could account for artifacts in proximity assays at endogenous membrane protein surface densities (Needham et al., 2013).

A higher density of ErbB2 proteins was observed in actin-GFP-rich membrane areas of the cell, such as peripheral ruffles, when compared to membrane areas with a low content of actin-GFP. This observation is in accordance with previous publications describing activated ErbB2 to be associated with actin-rich membrane protrusions in ErbB2 overexpressing cells (Benlimame et al., 2005; Brandt et al., 1999; Jeong et al., 2017; Jeong et al., 2016; Mori et al., 1987; Wang et al., 2006). The capability of our approach for single molecule detection led to the observation of QD-labeled ErbB2 homodimers in actin-GFP-rich peripheral membrane regions such as ruffles. The putative signaling active ErbB2 homodimers are thus preferentially found in membrane regions with an abundance of underlying actin.

An explanation for the spatial correlation of ErbB2 homodimers with peripheral membrane ruffles could be that ruffles resemble specialized regions that are not only abundant in densely packed actin filaments, but also in cholesterol (Grimmer et al., 2002; Qin et al., 2006) enriched lipid rafts (Balasubramanian et al., 2007). In fact, depletion of cholesterol inhibits the formation of membrane ruffling (Grimmer et al., 2002), and cholesterol enrichment induces ruffling in adherent cells (Qin et al., 2006). ErbB2 was indeed described to cluster within lipid rafts (**Figure 3C, D**) (Nagy et al., 2002). A high number of membrane ruffles is known to be associated with increased growth and invasive behavior of cancer cells (Jiang, 1995). Based on our finding of the direct relationship between actin-containing ruffles and signaling active ErbB2 homodimers, it can be speculated that the combined inhibition of the actin network and ErbB2 dimers resembles a promising therapeutic approach in cancer therapy.

In contrast to the observation of a peak at 20 nm, a peak at 36 nm was observed in the pair correlation function at peripheral membrane regions with a low content of actin-GFP. A protein distance of ~36 nm was formerly proposed to be caused by the association of ErbB2 with actin filaments (Parker et al., 2018). Direct binding of ErbB2 to actin filaments is unlikely because in contrast to EGFR, ErbB2 does not contain an actin binding domain (den Hartigh et al., 1992). The fact that the 36-nm peak was nevertheless observed, suggests that this inter-label distance is due to formation of EGFR:ErbB2 heterodimers, and the binding of EGFR to cortical actin, which is actin just below the plasma membrane. Others measured inter-label distances of fluorophore-labeled EGFR-Affibody by fluorophore localization imaging with photo bleaching (FLIP), and measured a value of 37 nm, which is of a similar dimension to the repeat of the left-handed helix of cortical, filamentous actin of 35.9 nm (Needham et al., 2014). The presence of a 36-nm peak in flat regions was confirmed in a control experiment in which membrane ruffles were destroyed via Cytochalasin D, producing a mostly flat-like membrane (**Figure 4**). However, the 36 nm peak was absent from data obtained for cells that were negative for actin-GFP in the fluorescence images (**Supplementary Figure S2**). It can thus be concluded that the hypothesized heterodimers become sufficiently abundant for observation only in cells with overexpressed and intact networks formed by actin-GFP.

We also expected a heterodimerization and thus a 36 nm peak for actin-rich peripheral ruffles. One reason for the absence of the 36 nm peak for this region could be that the number of ErbB2 homodimers was probably so large that the 36 nm peak was not visible in the pair correlation function (**Figure 3a**). Therefore, data was additionally analyzed by conducting a search for linear QD-chains in all data sets (**Figure 5**). Indeed, we were now able to detect linear particle chains with an inter-label distance of 36 nm ± 15 nm consisting of 6 or more particles in actin-GFP-rich membrane regions. In contrast, long, linear QD-chains remained undetected in actin-GFP low membrane regions, likely due to the limited labeling efficiency. This resulted in a reduced probability of detecting six or more particles in one line. The labeling efficiency is unknown; though assuming a high labeling efficiency of 70% would produce only a 0.7^6^ = 12% probability of finding a linear chain of six. This suggests that there may not have been sufficient QD-labels in the flat areas for detecting lines.

Thus, our data suggests that EGFR:ErbB2 heterodimers reside both inside and outside actin-GFP-rich regions. An implication of the presence of heterodimers would be a reduction of the membrane diffusion coefficient of a fraction of ErbB2 that becomes immobilized to actin via EGFR. Note that such immobilization would not occur for ErbB3 as this receptor form heterodimers with ErbB2 instead of EGFR (Varadi et al., 2019).

## CONCLUSION

By applying correlative fluorescence microscopy and STEM to single, QD-labeled ErbB2 proteins in actin-GFP transduced breast cancer cells; it was possible to detect distinct distribution patterns of membrane ErbB2 in dependency of the underlying actin content. Putative signaling active ErbB2 homodimers were detected in membrane regions with densely packed actin filaments, but not in regions without actin-GFP. The observation of a peak in the label pair distance of 36 nm suggests an interaction of ErbB2 with cortical actin filaments, most probably via heterodimerization with EGFR.

## DATA AVAILABILITY STATEMENT

All datasets generated for this study are included in the article/SupplementaryMaterial.

## AUTHOR CONTRIBUTIONS

IND, FL, NdJ designed the research. All authors wrote the manuscript. IND, and PT performed data analysis. IND, ZM performed the experiments.

## ACKNOWLEDGEMENTS

We thank S. Keskin, and P. Kunnas for experimental help, T. Trampert for technical support, and E. Arzt for his support through INM.

## SUPPLEMENTARY MATERIAL

The Supplementary Material for this article can be found online at:

**Figure S1.** Light microscopy of Actin- and ErbB2 stained SKBR3 cell displayed in Figure 2 of the main text to indicate membrane ruffles. **(A)** Peripheral ruffles appear as darker grey structures at the cell edge in the direct interference contrast (DIC) image. **(B)** Actin-green fluorescent protein (GFP) gives a strong signal at the same positions. **(C)** Quantum dot (QD)-stained ErbB2 molecules overlay with actin-containing peripheral ruffles. **(D)** Overlay of the DIC, GFP and QD-chanel as displayed in A-C.

**Figure S2.** ErbB2 stained SKBR3 cell without actin-GFP. **(A)** Cropped fluorescence micrograph of SKBR3 breast cancer cells showing QD-labeled membrane ErbB2 in red. Cell marked with * was lost during EM preparation process. **(B)** Corresponding low magnification scanning transmission electron micrograph of graphene covered breast cancer cells taken at the same area, Magnification *M* = 1000×. **(C)** High resolution scanning transmission electron (STEM) micrographs of region marked in a-b. QD-labels appear as white dots, *M* = 100,000×. **(D)** Pair correlation function *g*(*r*) as function of pair distance *r* of QD-labeled ErbB2 in flat regions without actin-GFP.

**Figure S3.** Correlative fluorescence microscopy and STEM of whole breast cancer cells after treatment with Cytochalasin D. **(A-C)** Cropped fluorescence micrographs of SKBR3 breast cancer cells showing cellular actin-GFP in green, and QD-labeled membrane ErbB2 in red. Areas were both signals overlay appears yellow. Image A was taken at the baseline before treatment with Cytochalasin D (1h, 2μM) and staining of QDs. Image B was acquired immediately after Cytochalasin D treatment and labeling of ErbB2 with QDs. Image C was generated of maximum intensity overlays of deconvoluted z-stacks from the same spot as in A and B. Cells that disappeared during Cytochalasin D treatment and EM preparation process are marked with white asterisks in image A. **(D)** Corresponding low magnification scanning transmission electron micrograph of graphene covered breast cancer cells taken at the same area, *M* = 1000×. **(E, F)** High resolution scanning transmission electron micrographs of region marked in a-d. QD-labels appear as white dots in F. E: *M* = 30,000×, F: *M* = 100,000×.

## SCHEMES

**Scheme 1.** Pseudo-code for detection of linear QD chains.

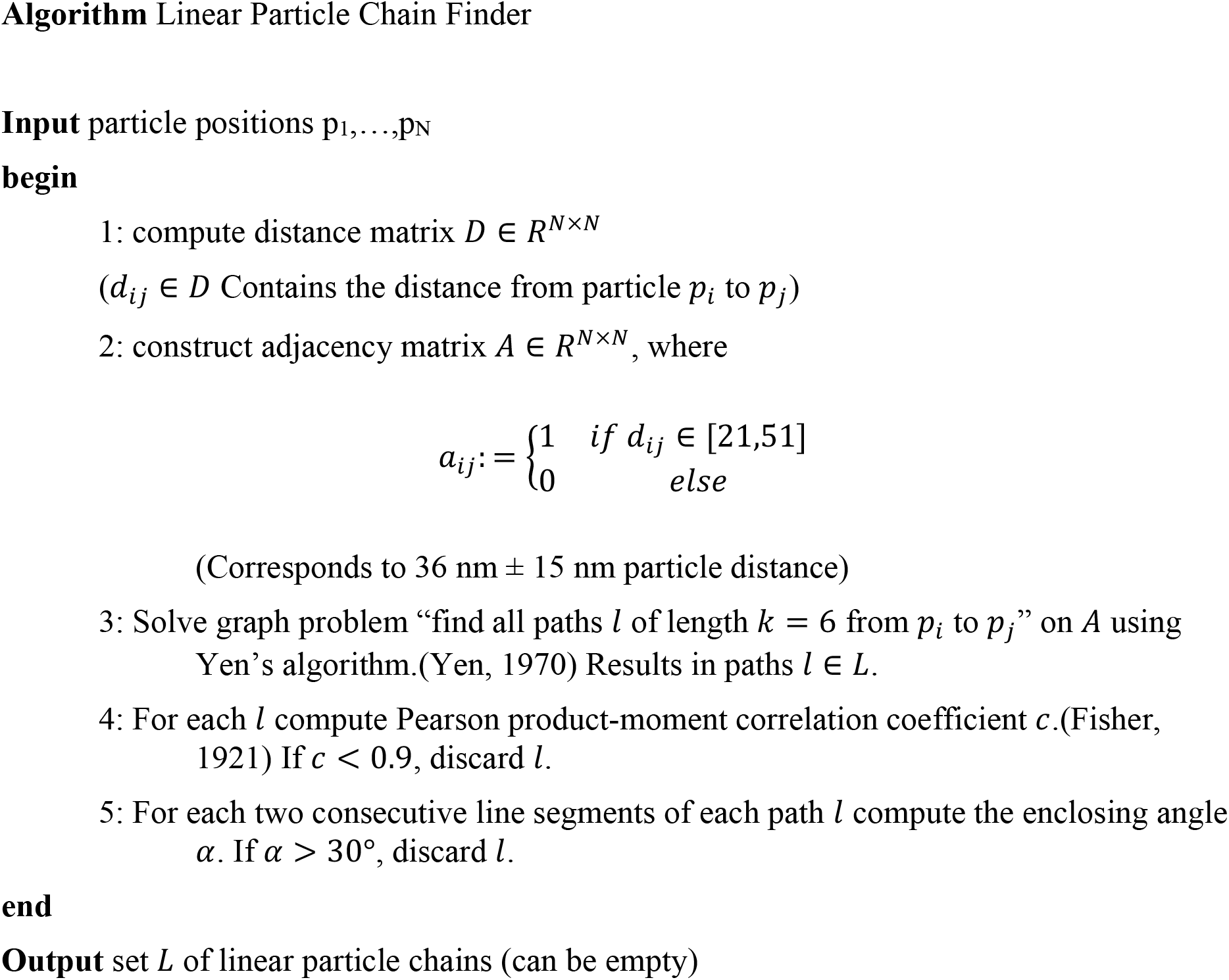

