## Supplementary Figures S1-S3 for "Correlative Fluorescence- and Electron Microscopy of Whole Breast Cancer Cells Reveals Different Distribution of ErbB2 Dependent on Underlying Actin"

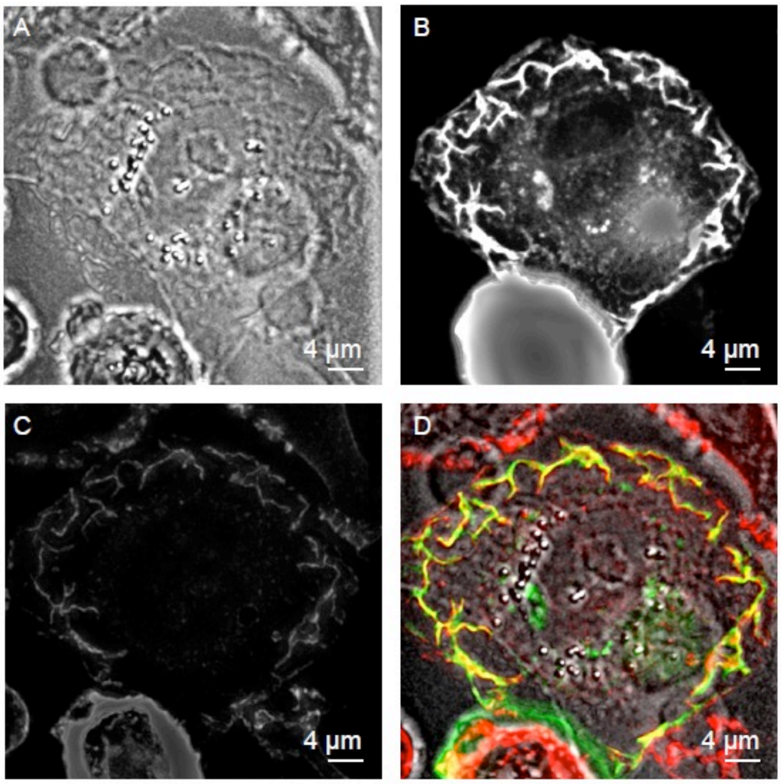


**Figure S1.** Light microscopy of Actin- and ErbB2 stained SKBR3 cell displayed in Figure 2 of the main text to indicate membrane ruffles. **(A)** Peripheral ruffles appear as darker grey structures at the cell edge in the direct interference contrast (DIC) image. **(B)** Actin-green fluorescent protein (GFP) gives a strong signal at the same positions. **(C)** Quantum dot (QD)-stained ErbB2 molecules overlay with actin-containing peripheral ruffles. **(D)** Overlay of the DIC, GFP and QD-chanel as displayed in A-C.

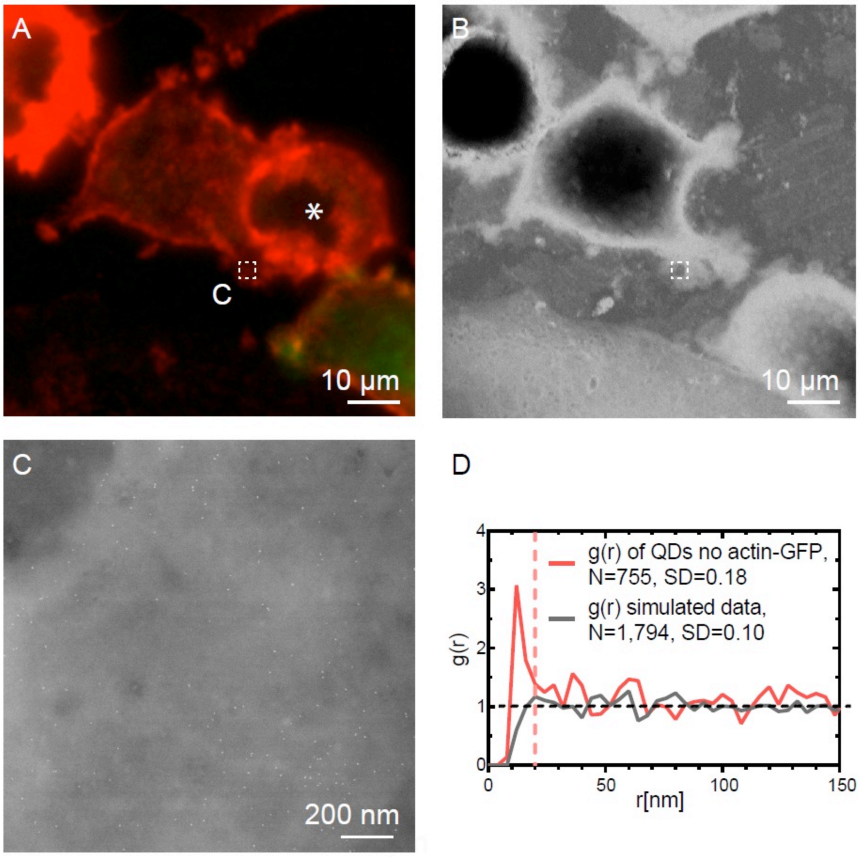


**Figure S2.** ErbB2 stained SKBR3 cell without actin-GFP. **(A)** Cropped fluorescence micrograph of SKBR3 breast cancer cells showing QD-labeled membrane ErbB2 in red. Cell marked with * was lost during EM preparation process. **(B)** Corresponding low magnification scanning transmission electron micrograph of graphene covered breast cancer cells taken at the same area, Magnification $M$ = 1000×. **(C)** High resolution scanning transmission electron (STEM) micrographs of region marked in a-b. QD-labels appear as white dots, $M$ = 100,000×. **(D)** Pair correlation function $g(r)$ as function of pair distance $r$ of QD-labeled ErbB2 in flat regions without actin-GFP.

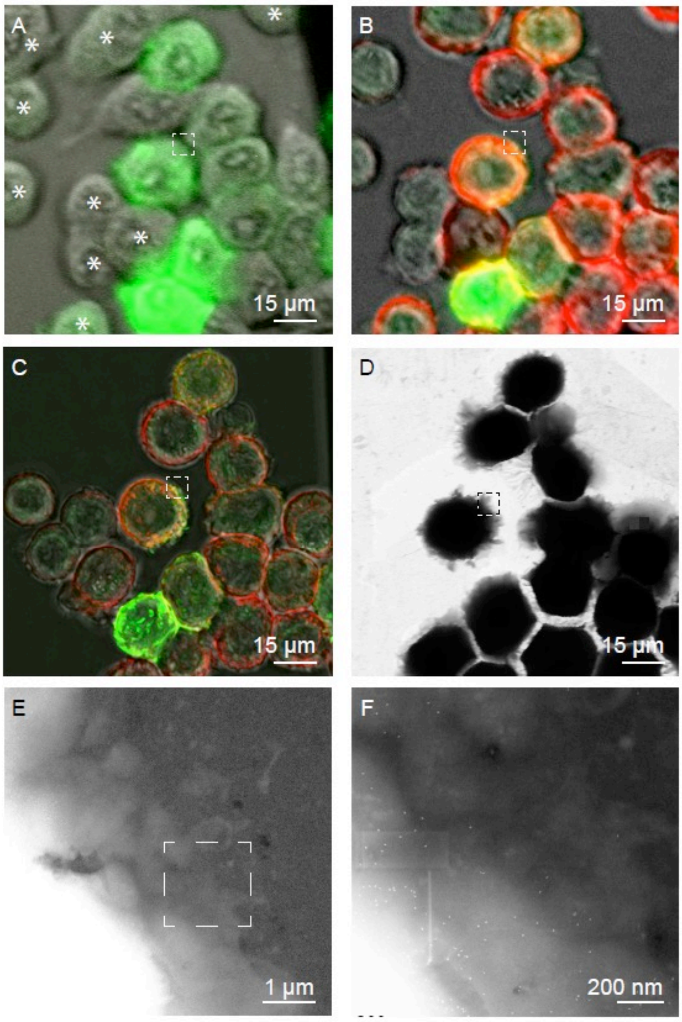


**Figure S3.** Correlative fluorescence microscopy and STEM of whole breast cancer cells after treatment with Cytochalasin D. **(A-C)** Cropped fluorescence micrographs of SKBR3 breast cancer cells showing cellular actin-GFP in green, and QD-labeled membrane ErbB2 in red. Areas were both signals overlay appears yellow. Image A was taken at the baseline before treatment with Cytochalasin D (1h, 2µM) and staining of QDs. Image B was acquired immediately after Cytochalasin D treatment and labeling of ErbB2 with QDs. Image C was generated of maximum intensity overlays of deconvoluted z-stacks from the same spot as in A and B. Cells that disappeared during Cytochalasin D treatment and EM preparation process are marked with white asterisks in image A. **(D)** Corresponding low magnification scanning transmission electron micrograph of graphene covered breast cancer cells taken at the same area, $M = 1000\times$. **(E, F)** High resolution scanning transmission electron micrographs of region marked in a-d. QD-labels appear as white dots in F. E: $M = 30{,}000\times$, F: $M = 100{,}000\times$.
